# Quantitative proteomic and metabolomic profiling reveals altered mitochondrial metabolism and folate biosynthesis pathways in the aging *Drosophila* eye

**DOI:** 10.1101/2021.06.02.446766

**Authors:** Hana Hall, Bruce R. Cooper, Guihong Qi, Aruna B. Wijeratne, Amber L. Mosley, Vikki M. Weake

## Abstract

Aging is associated with increased risk of ocular disease, suggesting that age-associated molecular changes in the eye increase its vulnerability to damage. Although there are common pathways involved in aging at an organismal level, different tissues and cell types exhibit specific changes in gene expression with advanced age. *Drosophila melanogaster* is an established model system for studying aging and neurodegenerative disease, that also provides a valuable model for studying age-associated ocular disease. Flies, like humans, exhibit decreased visual function and increased risk of retinal degeneration with age. Here, we profiled the aging proteome and metabolome of the *Drosophila* eye, and compared these data with age-associated transcriptomic changes from both eyes and photoreceptors to identify alterations in pathways that could lead to age-related phenotypes in the eye. Notably, the proteomic and metabolomic changes observed in the aging eye are distinct from those observed in the head or whole fly, suggesting that tissue-specific changes in protein abundance and metabolism occur in the aging fly. Our integration of the proteomic, metabolomic and transcriptomic data reveals that changes in metabolism, potentially due to decreases in availability of B vitamins, together with chronic activation of the immune response, may underpin many of the events observed in the aging *Drosophila* eye. We propose that targeting these pathways in the genetically tractable *Drosophila* system may help to identify potential neuroprotective approaches for neurodegenerative and age-related ocular diseases.

## Introduction

Age is the major risk factor for ocular disease, and is also a potential barrier to approaches aimed at regenerating cells in the eye (1–3). Thus, one of the most important questions in visual system biology is how increasing age predisposes the eye to disease. The eye is an oxygen-rich tissue with high metabolic needs due to the intensive energy requirements for light signaling, making it particularly vulnerable to defects in metabolism (4). In addition, maintenance of the proteome is critical to ocular health because many proteins within the light-sensing photoreceptor neurons in the eye must be continually regenerated to minimize accumulation of damage (5). Notably, while decreased proteostasis and mitochondrial dysfunction are hallmarks of aging across multiple tissues, defects in these processes have been shown to specifically increase the vulnerability of cells in the aging eye to disease (4, 6). Moreover, the eye is particularly sensitive to defects in metabolism that increase oxidative stress because of its high concentration of peroxidation-sensitive polyunsaturated fatty acids together with the high metabolic activity of the retina (4).

The fruit fly *Drosophila melanogaster* has been used as a model system for studying aging and neurodegenerative disease, providing insight into pathways that extend lifespan and protect against neuronal death (7, 8). *Drosophila* possess a compound eye that is composed of about 750 repeating units, termed ommatidia (Fig. 1A). Each ommatidium contains eight sensory neurons termed photoreceptors (also known as R cells: R1 – R8), four cone cells, and two primary pigment cells (9). Adjacent ommatidia share secondary and tertiary pigment cells, and mechanosensory bristles. The outer photoreceptors R1 – R6 express the light-sensitive G protein coupled receptor Rhodopsin 1 (Rh1, encoded by the *ninaE* gene) and are involved in motion detection, resembling vertebrate rods in function. In contrast, the inner photoreceptors R7 and R8 express other Rhodopsins sensitive to different wavelengths, with R7 stacked on top of R8 within each ommatidium. Each photoreceptor contains a light-sensitive organelle, the rhabdomere, that houses the phototransduction machinery. The eye is covered by a corneal lens, an extracellular secretion produced by the underlying cone and pigment cells. Below the corneal lens lies a gel-like substance termed the pseudocone, which together with the lens functions to focus light on the rhabdomeres within the retina (10). Photoreceptor axons project through the fenestrated membrane at the base of each ommatidium into optic ganglia in the brain (9).

**Fig 1.**
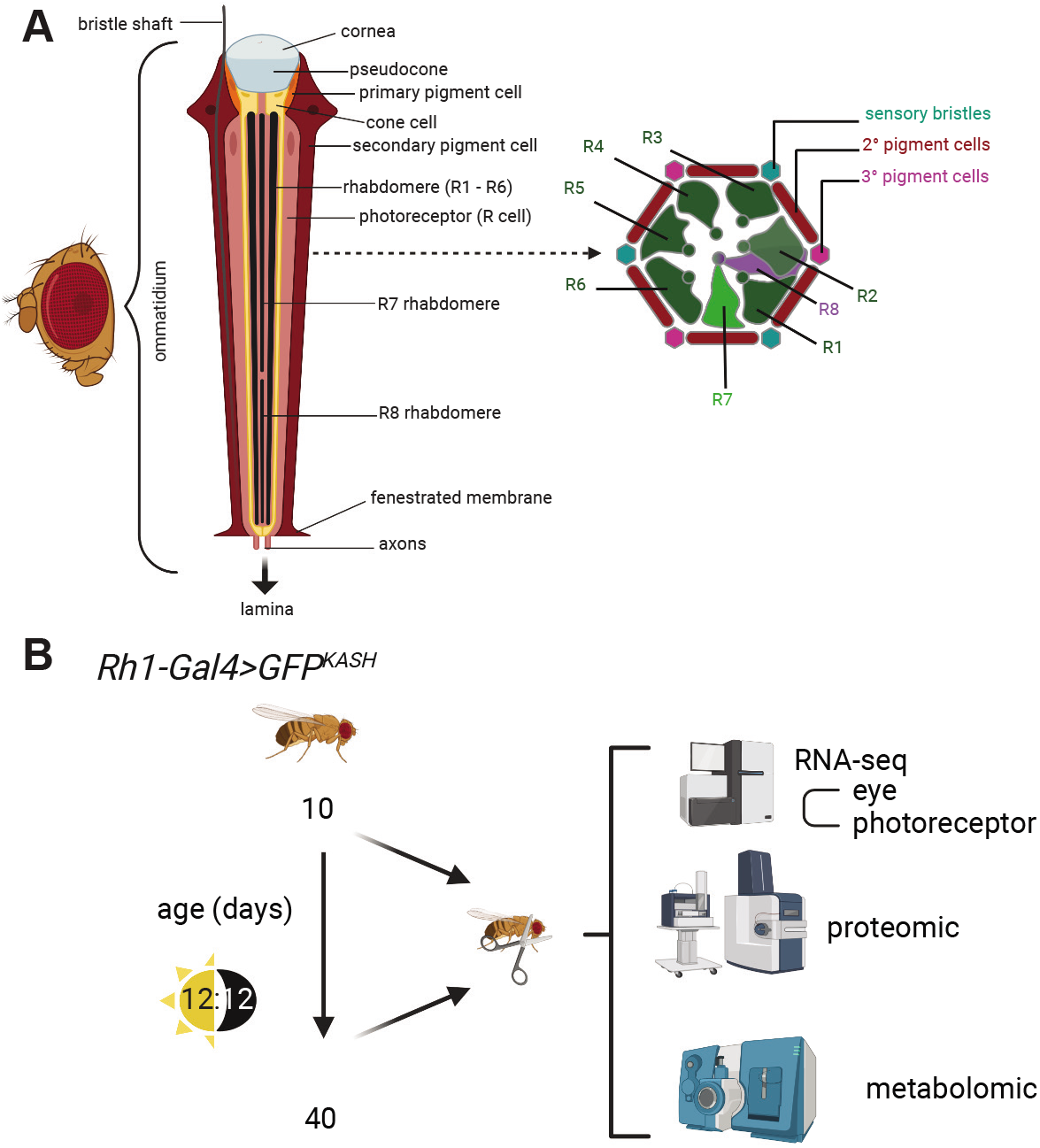
Comparison of the aging *Drosophila* eye transcriptome, proteome and metabolome. **(A)** The adult *Drosophila* compound eye is composed of approximately 700 ommatidia that contain photoreceptors, cone cells, pigment cells and a mechanosensory bristle. Section of an individual ommatidium (right) shows arrangement of outer photoreceptors (R1 – R6) and two inner photoreceptors (R7 and R8), which are stacked. Schematic adapted from (9). **(B)** Overview of the experimental scheme for quantitative analysis of the aging transcriptome, proteome and metabolome in *Drosophila* eye. Eyes were dissected from male *Rh1-Gal4>GFP^KASH^* flies at 10 or 40 days post-eclosion. Previously published RNA-seq data from eyes or photoreceptor nuclei (isolated using GFP^KASH^ tag) (11, 12) was compared with proteomic and metabolomic data obtained from this study.

As in vertebrates, *Drosophila* exhibit decreases in visual behavior with age that coincide with changes in gene expression and splicing, particularly in the light-sensing photoreceptor neurons (3, 11, 12). However, it is unclear whether these changes in gene expression are a cause or a consequence of other processes that become altered in the aging eye. To investigate this further, we sought to characterize the proteome and metabolome of the aging *Drosophila* eye and compare this with our previous characterization of the aging transcriptome of the eye and photoreceptors (11, 12). In addition, since recent studies have examined the aging proteome of *Drosophila* heads, containing both the brain and eye (14, 15), this provided the opportunity to ask whether the eye experiences unique proteomic changes that could explain its susceptibility to age-associated retinal degeneration.

## Results

### Global proteomic changes in the aging eye

We sought to characterize how age impacts protein abundance in the adult *Drosophila* eye. The dissected eyes used for proteomic analysis contain the lens, ommatidia, and lamina, together with a small amount of surrounding cuticle tissue, but do not contain additional tissues from the head or brain. To identify proteins with altered abundance in the aging eye, we performed a global quantitative comparison of eye proteins from adult male flies harvested at 10 or 40 days post-eclosion (hatching from the pupal case) using a tandem mass tags (TMT) “bottom-up” based quantitative proteomics mass spectrometry approach. We performed proteomic studies in male flies expressing a nuclear membrane-localized GFP tag in R1 – R6 photoreceptors, which facilitates photoreceptor-specific gene expression profiling: *Rh1-Gal4>GFP^KASH^* (11) (Fig. 1B). These flies have pigmented eyes and are otherwise wild type, exhibiting significant decreases in visual behavior between day 10 and day 40 without substantial retinal degeneration, similar to reports for standard wild-type fly strains such as OregonR (11). We used male flies for all analyses because there are sex-specific differences in visual behavior, eye size, and gene expression; male flies were also previously used for all aging RNA-seq studies in both photoreceptors and eyes (11, 12). We employed 10-plex TMT reagents to quantify the proteomes of eyes at day 10 (D10) or day 40 (D40), and identified 4351 proteins representing 4344 unique *Drosophila* protein isoforms (Fig. 2A, Table S1). 4046 of these protein isoforms were quantified in all four biological replicates from both ages, and were used for subsequent analysis. Principal component analysis (PCA) of the normalized protein abundances from each of the biological replicates revealed that 85% of the variance was associated with the difference in age between the sample groups, although smaller variation (PC2: 6%) was also present between samples at D10 but not in D40 (Fig. 2B). These data infer substantial changes in the eye proteome during aging, and suggest that increased age is associated with reduced heterogeneity in protein abundance in the eye.

**Fig 2.**
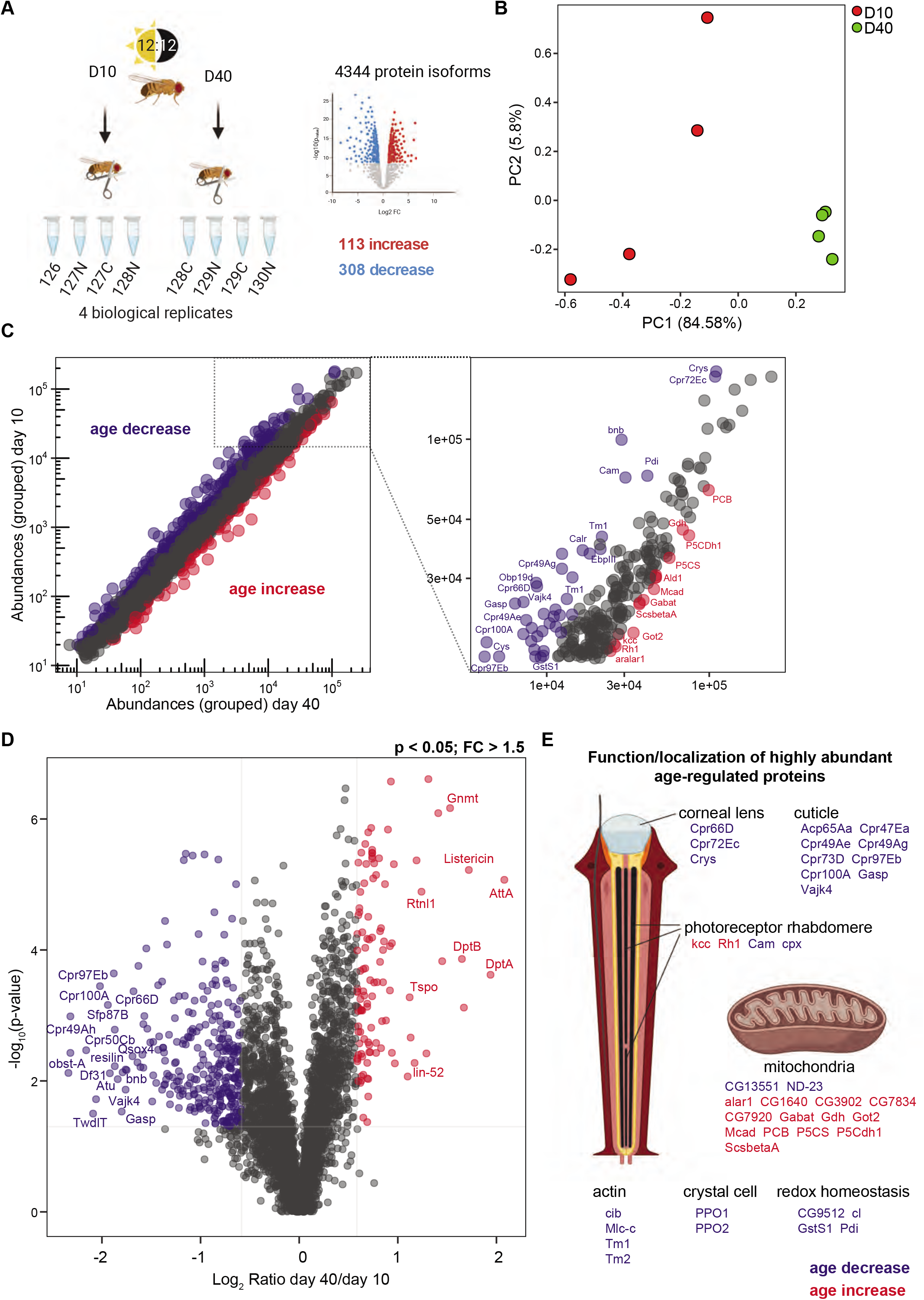
Quantitative proteomic analysis in the aging *Drosophila* eye. **(A)** Experimental scheme for quantitative analysis of the aging proteome in *Drosophila* eye. Eyes were dissected from male flies at 10 or 40 days post-eclosion, and quantitative proteomics employed using a “bottom-up” tandem mass tag multiplexed approach. Four biological replicates were analyzed for each age, with a total of 4351 proteins identified representing 4344 *Drosophila* protein isoforms. **(B)** Principal component analysis (PCA) of normalized protein abundance values for each sample at day 10 or 40. **(C)** Scatterplot displaying grouped protein abundances at day 10 versus day 40. Proteins that significantly decrease with increasing age are shown in blue, and proteins that significantly increase with age are shown in red (p < 0.05; fold-change >1.5). A magnified scatterplot in the right panel shows abundance ratios for a subset of highly abundant proteins (D10 grouped abundance > 15,000). Gene symbols for characterized *Drosophila* proteins are displayed for significantly altered proteins. **(D)** Volcano plot showing the log_2_ ratio of day 40/day 10 versus the −log_10_ p-value. Proteins with significant changes are highlighted in blue or red, as described in panel C. **(E)** The function and/or localization of highly abundant age-regulated proteins shown in panel C are displayed. Proteins that increase or decrease with age are shown in red or blue, respectively.

Next, we compared the abundance of proteins at D10 and D40 to identify proteins with abundance changes at each age using the criteria p < 0.05 and an abundance fold-change of 1.5 or greater). We identified 113 protein isoforms with significantly increased protein abundance at D40 relative to D10 (age increase) and 308 protein isoforms with significantly decreased protein abundance at D40 relative to D10 (age decrease). These changes in abundance were detected in proteins ranging over four orders of magnitude from the lowly expressed Regeneration (encoded by FBgn0261258 rgn, log_2_FC −0.8), to the highly abundant lens protein Crystallin (FBgn0005664, Crys; log_2_FC −0.7) (Fig. 2C). Almost three times as many proteins showed decreased rather than increased abundance with age (Fig. 2D), consistent with observations in other aging proteomic studies in *Drosophila* suggesting that there is a progressive reduction in protein synthesis during aging (14). We highlight the function of a subset of the most highly abundant proteins (Fig. 2C, right panel) that show altered abundance in the aging eye in Figure 2E, and the function of these proteins will be discussed further in a later section of the results.

### Tissue-specific changes in protein abundance occur in the aging eye

Other studies have examined the aging proteome of *Drosophila* heads and identified proteins that show significant changes in abundance with age (14, 15). Thus, we sought to compare our eye proteomic data with the available published data to determine if the eye exhibited specific changes during aging that were not detected in whole heads. We identified 4344 unique *Drosophila* protein isoforms in the eye, which is similar to the 4014 proteins identified in the head by Yang et al. using a similar TMT-multiplexing proteomics approach (14), and higher than the 1281 proteins identified in the head by Brown et al. using a different mass spectrometry approach (15). When we compared the proteins identified in the Yang et al. study in the head with the proteins we identified in the eye, we found that 3243 proteins were detected in both data sets, with 1099 proteins uniquely identified in the head and 861 proteins only identified in the eye (Fig. 3A, Table S2). While we expected that some proteins might be specifically expressed in the brain but not in the eye, and would therefore be unique to the head proteomic data set, we had expected all of the proteins detected in the eye also to be present in the head data. We assumed that some of the eye-specific proteins were not detected in the head proteomic analysis because they are expressed at low levels. To test this hypothesis, we examined the abundance of the 861 eye-specific proteins relative to the 3243 proteins that were detected in both the head and eye (shared proteins). We found the proteins detected only in the eye were substantially less abundant compared to the proteins detected in both tissues suggesting the presence of these lowly expressed proteins may be masked in proteomic data from the entire head (Fig. 3B). Interestingly, when we performed the analysis in the other direction, we found that the 1099 proteins that were only detected in the head were also less abundantly expressed compared with the shared proteins (Fig. 3C). We conclude that these proteins are likely expressed specifically within the brain, and they are either present at very low levels or are entirely absent in the eye.

**Fig 3.**
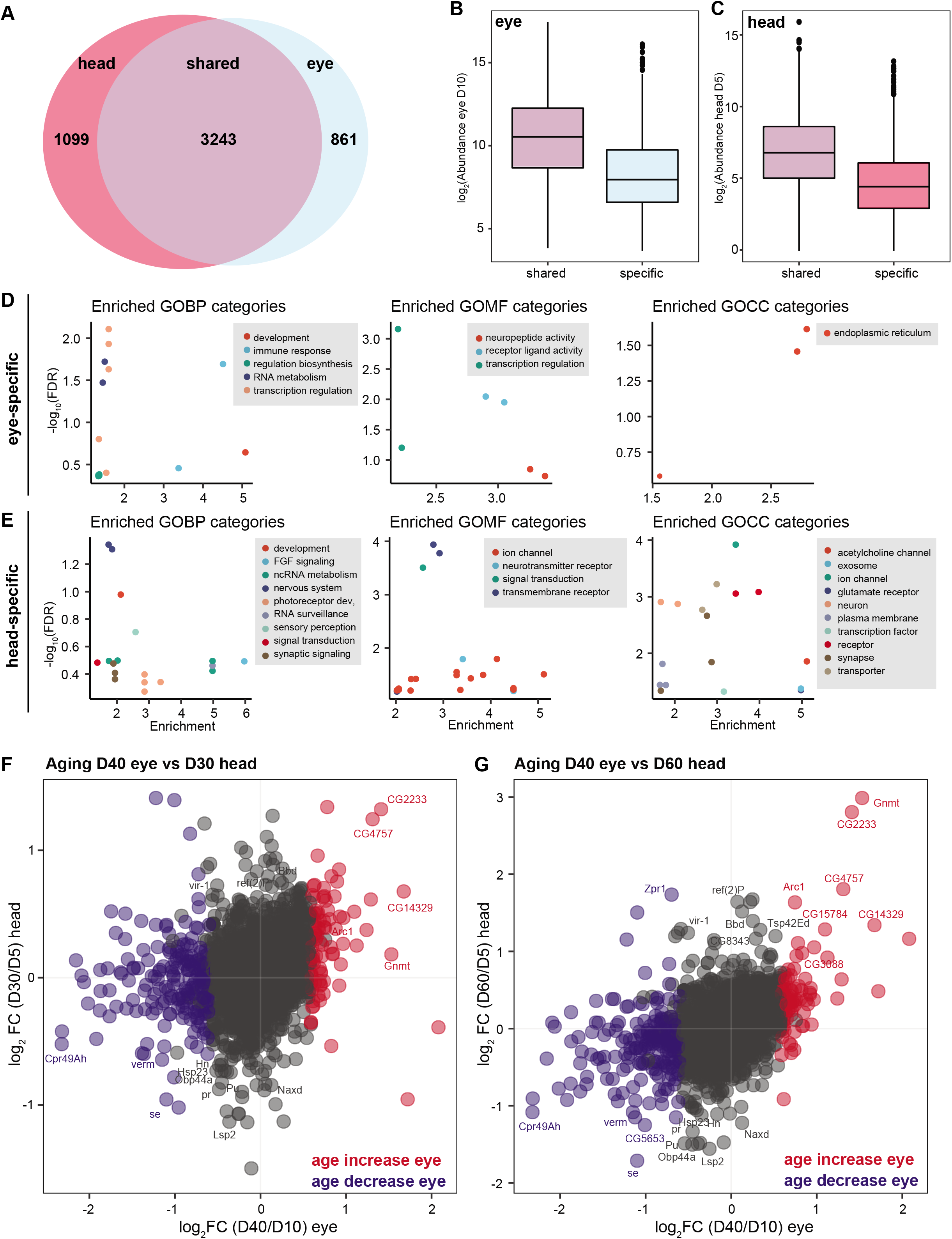
Comparison of the aging eye and head proteome. **(A)** Venn diagram showing the overlapping (shared) and unique protein isoforms identified in the eye in this study compared with published data from heads. **(B)** Boxplot showing protein abundance in eyes from young flies (log_2_ abundance D10) for proteins detected in both eyes and heads (shared) or only in eyes (specific). **(C)** Boxplot showing protein abundance in heads from young flies (log_2_ abundance D5) for proteins detected in both eyes and heads (shared) or only in heads (specific). **(D, E)** Enriched GO biological process (GOBP), molecular function (GOMF) and cellular compartment (GOCC) categories for 861 proteins uniquely detected in eyes (panel D) or 1099 proteins uniquely detected in heads (panel E) compared to all proteins detected in both data sets. **(F, G)** Scatterplots showing the log_2_ ratio of day40/day 10 for proteins in the eye versus head at day 30/day 5 (F) or day 60/day 5 (G). Color indicates proteins that significantly increase (red) or decrease (blue) with age in the eye. The gene symbol is shown for a subset of proteins with significant changes in the aging head.

Next, we raised the question if the eye- or head-specific proteins had functions that could be unique to either tissue. To answer, we performed Gene Ontology (GO) term analysis on the eye- or head-specific proteins relative to all proteins identified in either proteomic data set. The eye-specific proteins were enriched for biological processes associated with transcription regulation and immune response, and for the endoplasmic reticulum as a cellular component (Fig. 3D, Table S3). In contrast, GO molecular functions were highly enriched for ion channels and receptors in the head-specific proteins (Fig. 3E, Table S3). Surprisingly, the head-specific proteins were also enriched for GO terms associated with photoreceptor cell fate commitment. These head-specific proteins include several proteins involved in specification of the R7 photoreceptor cell during eye development: Sevenless (sev; FBgn0003366) and Son of sevenless (Sos; FBgn0001965). Although Sevenless signaling plays a key role in eye development, it is expressed only transiently in the eye imaginal disc during the larval stage, and is not detected in pupae (16). However, Sevenless is also expressed in specific neurons in the larval central nervous system (17) and is detected in adult heads (16), potentially explaining the unexpected identification of this protein in adult heads, but not in eyes.

Last, we compared the changes in abundance in the shared 3243 proteins during aging in the eye and head. Whereas we examined protein abundance at D10 and D40, Yang et al. compared D5 with either D30 or D60 (14). Thus, we compared the changes in protein abundance in the eye between D10 and D40 with changes in the head at D30 relative to D5 (Fig. 3F) or D60 relative to D5 (Fig. 3G). Overall, the age-related changes in protein abundance in the eye showed a similar trend to those in the head from older flies at D60, with a much weaker correlation to younger flies at D30 (compare Fig. 3F and G). Moreover, only 60 proteins were differentially regulated with age in both the older D60 heads and eyes, and several of these show changes in the opposing direction (Fig. 3G). Together, these data indicate that there are substantial differences in aging within the eye relative to the brain and the remainder of the head tissues. Moreover, the eye appears to experience age-related changes in protein abundance much earlier than in the brain, potentially due to its exposure to damaging light and the high metabolic activity associated with visual perception.

### Mitochondrial enzymes involved in metabolism are upregulated in the aging eye

Since changes in the abundance of highly expressed proteins might provide insight into the biological outcome of aging in the eye, we first examined the function of a subset of the most highly abundant proteins that were differentially regulated with age (Fig. 2C, right panel). We observed a substantial number of mitochondrial proteins in the differentially expressed highly abundant proteins, the majority of which showed increased abundance with age (Fig. 2E).

These include enzymes involved in the tricarboxylic acid (TCA) cycle such as Succinyl-coenzyme A synthetase β subunit, ADP-forming (ScsbetaA; FBgn0037643) and enzymes involved in glutamate metabolism such as Glutamate dehydrogenase (Gdh; FBgn0001098), Delta[1]-pyrroline-5-carboxylate synthase (P5CS; FBgn0037146), delta-1-Pyrroline-5-carboxylate dehydrogenase 1 (P5CDh1; FBgn0037138), and the mitochondrial glutamate transporter Glutamate Carrier 1 (GC1; FBgn0260743). GO term analysis of all 113 age increased proteins relative to all detected proteins also revealed metabolic processes including amino acid metabolism, glutamine/glutamate metabolism, and proline metabolism as being highly enriched (Fig. 4A, Table S4). We also identified the mitochondrion as the sole enriched GO cellular component in the age increased proteins (Fig. 4A), suggesting that mitochondrial enzymes involved in metabolism show an overall increase in abundance in the aging eye.

**Fig 4.**
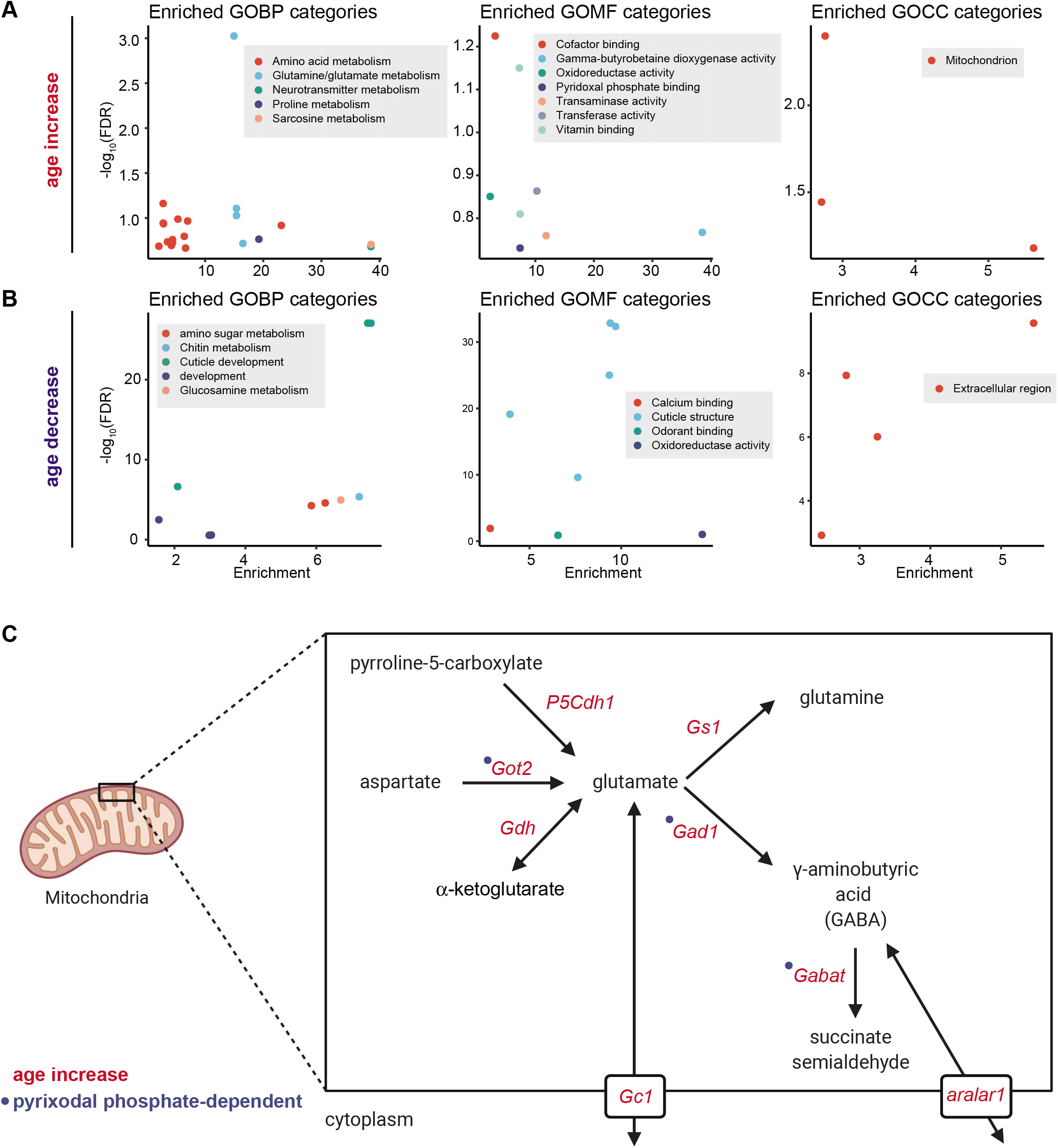
Functional analysis of significantly altered proteins in the aging *Drosophila* eye. **(A, B)** Enriched GO biological process (GOBP), molecular function (GOMF) and cellular compartment (GOCC) categories for 113 proteins that increase (panel A) or 308 proteins that decrease (panel B) with age compared to all 4344 proteins detected in adult eyes. **(C)** Enzymes and transporters involved in glutamate and GABA metabolism show increased levels in the aging eye. Proteins with significant increases during aging are highlighted in red, and enzymes that bind pyridoxal phosphate are marked by blue circles. The subcellular localization of enzymes and transporters is shown for illustrative purposes within a single cell, but these proteins may be expressed specifically in different cell types within the eye.

When we closely examined the function of the metabolic enzymes that showed increased abundance with age, we found that several of the enzymes involved in amino acid metabolism and glutamine/glutamate metabolism are also key regulators of neurotransmitter levels in the fly brain. Notably, we identified neurotransmitter metabolism as one of the enriched GO terms in the age increased proteins (Fig. 4A, Table S4). Glutamate functions as a neurotransmitter, and lamina neurons involved in motion vision express the sole vesicular glutamate transporter in flies, VGlut (FBgn0031424) (18). All three of the glutamate-metabolizing enzymes that regulate glutamate neurotransmitter pools showed increased abundance with age including Glutamate oxaloacetate transaminase 2 (Got2; FBgn0001125), Glutamic acid decarboxylase 1 (Gad1; FBgn0004516) and Glutamine synthetase 1 (Gs1; FBgn0001142) (19) (Fig. 3C). Gad1 is necessary for synthesis of another neurotransmitter, γ-aminobutyric acid (GABA). There is also an age-associated increase in abundance of the mitochondrial transporter aralar1 (FBgn0028646) that has been shown to sequester GABA in response to increased mitochondrial activity (20), as well as increased levels of one of the major enzymes involved in GABA catabolism: Gamma-aminobutyric acid transaminase (Gabat, FBgn0036927). In addition to glutamate and GABA, other enzymes that affect levels of neurotransmitters or neuromodulators are altered during aging in the eye. For example, Pyruvate carboxylase (PCB; FBgn0027580), which has roles in replenishing key TCA intermediates, also negatively affects memory formation by decreasing levels of serine, which functions as a neuromodulator (21). We observed increased PCB levels in the aging eye, and other studies have shown that PCB activity increases in *Drosophila* glial cells during aging (21). Together, these data suggest that there are alterations in the pathways involved in neurotransmitter metabolism in the aging eye that could impact visual behavior, particularly in lamina neurons within the optic ganglia.

### The aging eye shows decreases in abundance of cuticular proteins including constituents of the corneal lens

When we examined the most abundant proteins that showed decreased levels during aging, it was immediately apparent that cuticular proteins (Cpr genes) were highly enriched in this group (Fig. 2C, E). The fly eye is surrounded by a cuticle that is composed of chitin and cuticular proteins, some of which are also components of the corneal lens (22). Three of the four characterized protein components of the corneal lens showed significant decreased abundance during aging: Cpr66D (FBgn0052029), Cpr72Ec (FBgn0036619), and Crys (22). The last of the four known corneal lens proteins, retinin (FBgn0040074), was detected in the proteomic analysis but was not significantly altered with age (Table S1). GO term analysis of all 308 age decreased proteins relative to all detected proteins revealed chitin metabolism and cuticle development as highly enriched biological processes (Fig. 4B, Table S4). Moreover, the only enriched GO cellular component terms for the age decreased proteins was the *extracellular region*, suggesting that the aging *Drosophila* eye shows substantial decreases in the levels of many proteins associated with extracellular structures such as the corneal lens. These changes may affect the composition and structure of the lens, potentially altering its ability to focus light on the apical surface of photoreceptors thereby affecting visual function.

### Proteins involved in calcium buffering and redox homeostasis show decreased levels in the aging eye

When we examined the GO molecular function enrichment for proteins that showed decreased levels with age, we observed an enrichment for calcium binding and oxidoreductase activity. In the presence of light, activation of the phototransduction cascade by the G-protein coupled receptor Rhodopsin results in opening of the transient receptor potential (trp) channels and influx of calcium into photoreceptors (23). Prolonged high levels of intracellular calcium can be toxic, and calcium is rapidly extruded by exchangers such as Calx (23). We observed a more than two-fold decrease in levels of the calcium binding protein Calmodulin (Cam, FBgn0000253) in the aging eye (Fig. 2C, E; Table S1). Calmodulin is required for calcium-dependent deactivation of Rhodopsin, and the rapid termination of the light response is extremely sensitive to levels of this protein (24). Rhodopsin 1 (Rh1, ninaE; FBgn0002940) levels increased with age (Fig. 2C, E), suggesting that the phototransduction cascade can be activated to the same, or potentially higher, levels in photoreceptors from older flies. Consistent with this, analysis of electroretinograms from pigmented (w^+^) flies similar to the genotype used in our study showed that the receptor potential, which represents photoreceptor activation, does not differ substantially between young and old flies (25). In addition to Calmodulin, we observed a substantial decrease in age-associated levels of the calcium buffering protein Calphotin (Cpn, FBgn0261714), which protects photoreceptors from light-induced retinal degeneration (26). Like Calmodulin, photoreceptors are sensitive to levels of Cpn, suggesting that the two-fold decrease in levels of Cpn in the aging eye might reduce the ability of photoreceptors to buffer the increase in intracellular calcium concentration in the light, enhancing the risk of retinal degeneration. Previous studies in our lab have shown that calcium influx in response to an acute light stress induced by blue light exposure results in lipid peroxidation, causing retinal degeneration (27). Intriguingly, we observed a significant decrease in levels of Glutathione S transferase S1 (GstS1; FBgn0010226), which specifically conjugates the lipid peroxidation product 4-hydroxynonenal (4-HNE) (28). Overexpression of GstS1 has been shown to suppress neurodegeneration in *Drosophila* models of Parkinson’s disease (29); thus, even modest decreases in the levels of GstS1 may increase levels of toxic lipid peroxidation products in the aging eye.

### Proteome changes in the aging eye reflect transcriptome changes in multiple cell types in the eye including photoreceptors

Since the eye is composed of multiple cell types (Fig. 1A), we next compared the proteomic changes observed in the aging eye with transcriptomic changes previously examined either in the whole eye (12) or in enriched photoreceptor nuclei (11). As mentioned previously, all studies were performed in male flies of the same genotype to facilitate direct comparisons between each data set (Fig. 1B). The 4335 *Drosophila* proteins that were identified in our proteomic analysis mapped to 3863 transcripts from unique genes expressed at detectable levels in the eye, and 3388 transcripts expressed in R1 – R6 photoreceptors (Fig. 5A). Because more than half of the genes detected by RNA-seq analysis were not detected in the proteomic data set, we first asked if these undetected proteins corresponded to less abundantly expressed genes. Indeed, when we examined the expression of transcripts (RPKM, Reads Per Kilobase of transcript per Million mapped reads) encoding proteins that were not detected in our proteomics analysis (Fig. 5B, not detected, ND), these genes showed much lower relative expression as compared with more abundant proteins. Further, relative protein abundance correlated with RNA expression more strongly in the eye than in photoreceptors, although overall transcript expression was higher for the most abundant proteins (Q4) relative to least abundant proteins (Q1) in both the eye and photoreceptors (Fig. 5B). Moreover, more than a quarter of the least abundant proteins were not detected at the transcript level in photoreceptors (Fig. 5B, Q1), suggesting that these proteins might be expressed in other cell types in the eye such as the cone, pigment, or bristle cells.

**Fig 5.**
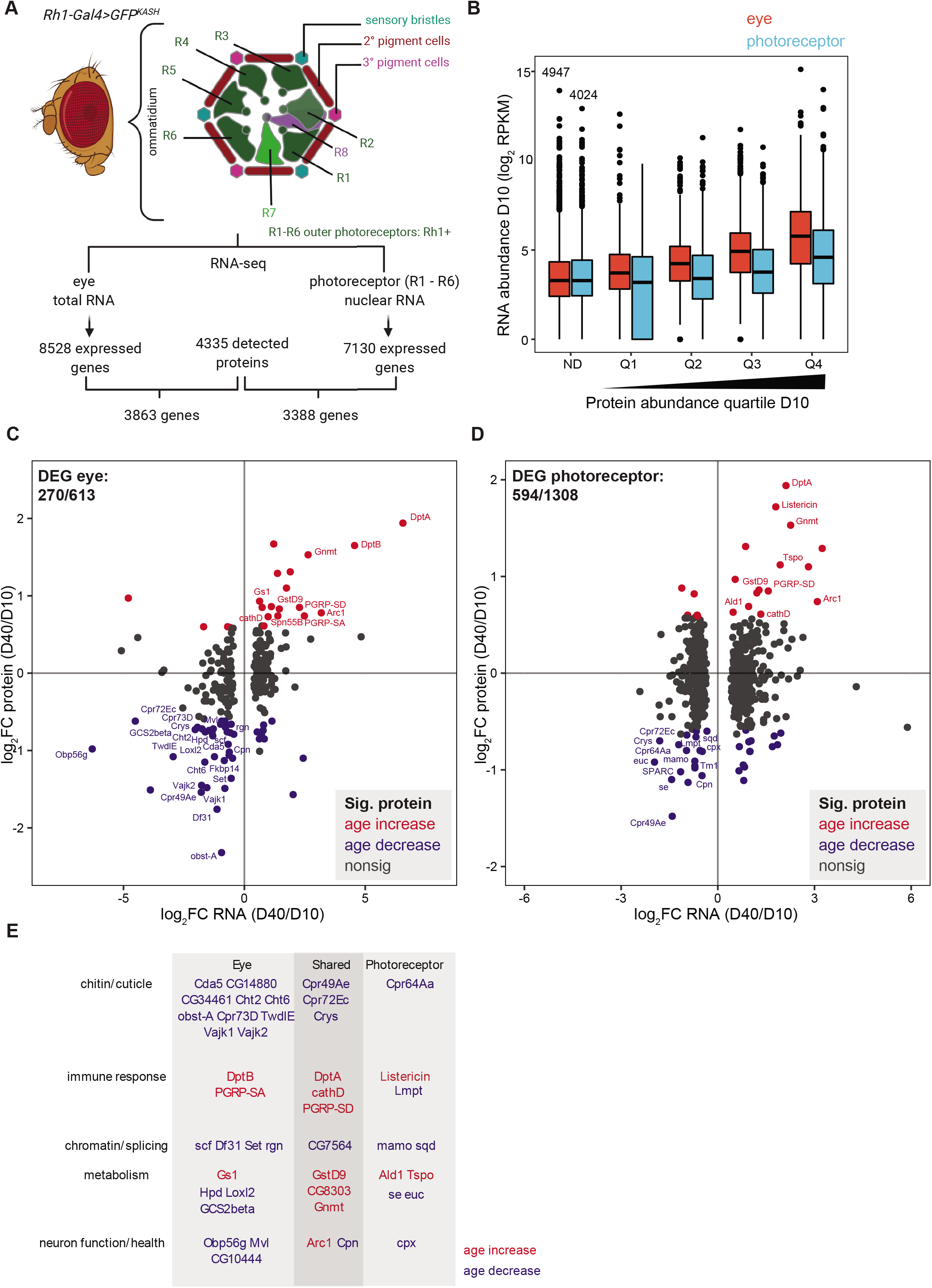
Proteome changes in the aging eye reflect transcriptome changes in multiple cell types in the eye including photoreceptors. **(A)** Proteins identified in the aging eye were compared with expressed genes previously detected in the eye or photoreceptors by RNA-seq. The *Drosophila* compound eye is composed of ommatidia that contain different cell types including the outer R1 – R6 photoreceptors, which are labeled with *Rh1-Gal4>GFP^KASH^* for photoreceptor nuclear RNA-seq. **(B)** Boxplot showing RNA abundance (log_2_ RPKM D10) for all expressed genes in eyes or photoreceptors. Proteins were separated into expression quartiles based on relative abundance at D10 (Q1 low, Q4 high; ND, protein not detected). One was added to RPKM values of 0; these appear on the log_2_ scale as 0, representing genes that were not detected in RNA-seq data. The number of expressed genes (RNA) not detected at the protein level is shown above the boxplot for ND. **(C,D)** Scatterplots showing the log_2_ ratio of day 40/day 10 for RNA versus protein in eye (C) or photoreceptor (D). The number of differentially expressed genes (DEGs; FDR < 0.1) in RNA-seq analysis that were present in the proteomic analysis is shown in the upper left quadrant. Color indicates proteins that significantly increase (red) or decrease (blue) with age. The gene symbol is shown for characterized genes that change in the same direction in both proteomic and transcriptomic analysis. **(E)** Function of selected proteins with altered expression at both the RNA and protein level in the eye, photoreceptor, or in both comparisons (shared). Color indicates proteins that significantly increase (red) or decrease (blue) with age.

Next, we compared all the differentially expressed genes between day 10 and day 40 that were previously identified using DESEQ2 analysis of RNA-seq data in the eye (613 genes; FDR < 0.01) or photoreceptors (1308 genes; FDR < 0.01) with the proteins that were detected in our proteomic analysis (12). We detected 270 of the 613 differentially expressed genes in the eye in the proteomic analysis (DEGs; Fig. 5C), 53 of which showed significant expression changes in the same direction at both the transcript and protein levels. However, most of the DEGs in the eye also showed similar directions of change in expression at the protein level, even if these were not significant (Fig. 5C). In contrast, while 594 of the 1308 DEGs in photoreceptors were also present in the proteomic analysis, only 33 were significantly regulated in the same direction, and many DEGs showed opposing directions of change at the protein versus transcript level (Fig. 5D). Thus, we conclude that the overall changes in protein abundance in the eye reflect contributions from multiple cell types that likely mask some of the changes within individual cell types in the eye such as photoreceptors. Moreover, differences between the aging transcriptome and proteome in the eye may reflect changes in protein synthesis or post-translational abundance rather than transcript abundance alone. Despite this, we observed some similarities in the overlapping functional categories between the genes that were significantly regulated in the same direction using both approaches. For example, many of the transcripts for cuticle proteins were significantly downregulated with age, particularly in the eye (Fig. 5E), matching the decrease in levels of cuticle proteins with age (Fig. 2E, 4B). The lens is secreted during pupal development (9), and thus the decrease in levels of cuticular proteins may represent the diminishing expression of these proteins in adult flies following the end of development. Consistent with this idea, we previously observed a significant decrease in expression of many of these same cuticular proteins in photoreceptors in the first few days following eclosion (13). Similarly, we observed an increase in expression of genes involved in innate immunity such as Diptericin A (DptA; FBgn0004240), Diptericin B (DptB; FBgn0034407), and Listericin (FBgn0033593) in the first few days after eclosion (13), suggesting that these proteins may be induced in the adult eye shortly after development is completed, with continued increases in expression throughout the adult lifespan. We also observed increased expression of several peptidoglycan recognition proteins (PGRPs) that are known to upregulate the immune response at both the transcript and protein level (30), suggestive of a chronic activation of the immune response in the aging eye. Notably, chronic activation of the immune response due to constitutive expression of PGRP results in reduced lifespan in flies (31). Activation of the immune response may also result in increases in expression of some metabolic enzymes such as Glycine N-methyltransferase (Gnmt, FBgn0038074) and the outer mitochondrial membrane Translocator protein (Tspo, FBgn0031263) (Fig. 5E). Inflammation has been shown to induce Gnmt expression in fat body (32), and Tspo knockdown results in defective responses to infection (33). Intriguingly, Gnmt and Tspo have opposing effects on longevity in flies: overexpression of Gnmt extends lifespan (34), whereas inhibition of Tspo prolongs lifespan and protects against neurodegeneration (35). These data suggest that chronic activation of the immune response in the aging eye occurs primarily at the transcriptional level, and may impact metabolism and mitochondrial function, particularly in photoreceptors that have high metabolic requirements due to the intensive energy needs necessary for light sensing.

### Metabolomic changes in the aging eye impact pterin biosynthesis and could alter protein methylation

To examine if aging caused global changes in metabolism in the eye, we used an unbiased mass spectrometry approach to profile the metabolome of eyes from flies at day 10 and day 40 (Fig. 6A). Using this approach, we identified 1270 unique compounds with specific m/z ratios and retention times, 1148 of which were identified in all four of the biological replicates in at least one of the age groups. PCA analysis of the normalized compound abundances demonstrated that 66% of the variance was associated with age, while a smaller difference existed between samples particularly at day 40 (PC2: 12%) (Fig. 6B). This suggests that older eyes have increased heterogeneity in metabolite abundance relative to young eyes, a finding that directly contrasts with the observations from the aging proteomic studies (compare Fig. 2B and Fig. 6B). We next compared day 10 and day 40 metabolite profiles to identify compounds that were significantly changed with age in the eye (p < 0.01; fold-change >2). Using this approach, we identified 103 compounds that increased with age, and 168 compounds that decreased with age (Fig. 6A, C, Table S5). To identify these compounds, we first searched the METLIN and BioCyc databases for matches based on exact mass (Table S6). Since many compounds had potential matches to multiple metabolites, we also performed HPLC-MS/MS to generate fragmentation spectra from pooled day 10 and day 40 samples. The resulting spectra and retention times were used to identify a subset of the significantly altered compounds with high confidence (Fig. 6C, Table S6). Last, we used mummichog to identify metabolic pathways that were enriched for all of the altered metabolites identified in the aging eye. Using this approach, we observed strong enrichment for metabolites associated with folate biosynthesis and related pathways including one carbon pool by folate, porphyrin metabolism, and purine metabolism (Fig. 6D, Table S7). Most of the metabolites associated with the folate pathway showed decreased levels in the aging eye including neopterin, 7,8-dihydro-biopterin, sepiapterin, isoxanthopterin, and biopterin. In addition, levels of purines such as guanosine, guanine, adenine and adenosine were also decreased in the aging eye, together with phenylalanine (Fig. 6D, Table S6 and S7). Interestingly, we observed a significant decrease in the levels of riboflavin (Vitamin B2) in the aging eye (Fig. 6C, Table S6). We confirmed the identity of riboflavin by comparison of the retention time and MS/MS spectra, as well as using commercially available standards. Riboflavin is a precursor for the synthesis of Flavin mononucleotide (FMN) and Flavin adenine dinucleotide (FAD), which are cofactors for enzymes involved in many different redox reactions (36). Thus, it is possible that decreased riboflavin levels in the aging eye might impact the activity of many different metabolic pathways.

**Fig 6.**
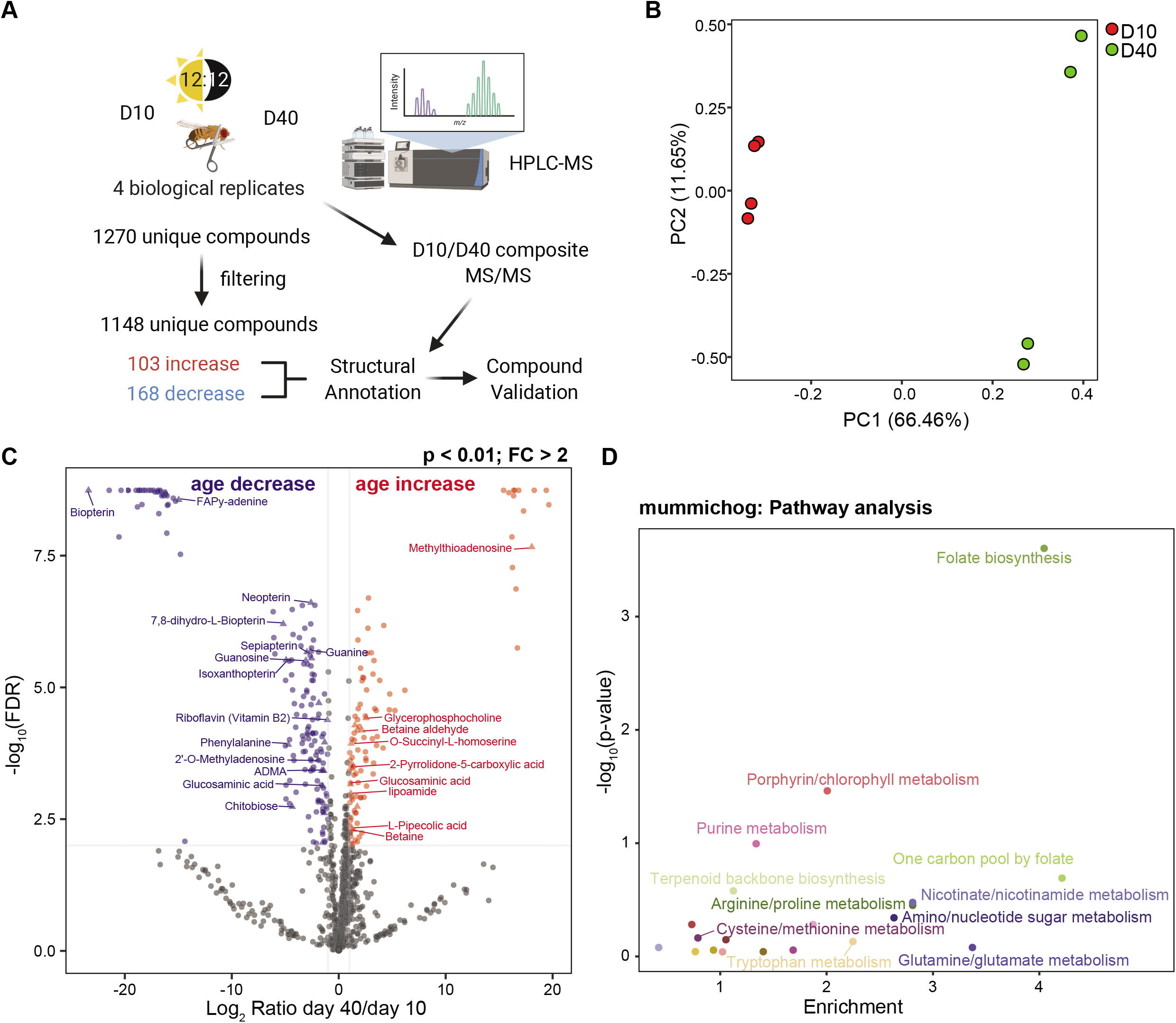
Global metabolome changes in the aging eye affect folate and one carbon metabolism pathways. **(A)** Experimental scheme for global metabolomic analysis of the aging *Drosophila* eye. Eyes were dissected from male flies at 10 or 40 days post-eclosion, and non-targeted metabolite profiling was performed on methanol-extracted samples using HPLC-MS. Unique compounds were filtered and normalized to identify 1148 unique compounds detected in all four biological replicates for either age. 271 compounds with significantly different levels at day 40 versus day 10 (p < 0.01; fold-change >2) were annotated using the METLIN database. The identify of a subset of compounds was validated by performing MS/MS on composite libraries prepared from each age, and using commercial standards. **(B)** Principal component analysis (PCA) of normalized metabolite abundance for each sample at day 10 or 40. **(C)** Volcano plot showing the log_2_ ratio of day 40/day 10 versus the −log_10_ p-value. Compounds that significantly decrease with increasing age are shown in blue, and compounds that significantly increase with age are shown in red (p < 0.01; fold-change >2). The names of a subset of compounds that were identified by MS/MS and/or standards are shown. **(D)** Mummichog pathway analysis was performed for the 271 age-regulated compounds. Scatterplot displaying enrichment versus −log_10_ p-value for identified pathways.

In contrast to the decrease in pterin derivatives, we observed significant increases in several metabolites involved in cysteine/methionine metabolism. Several of these metabolites are involved in regeneration of the methyl donor S-adenosyl-methionine (SAM), which is required for methylation reactions. For example, betaine and betaine aldehyde, which can be used to regenerate methionine from homocysteine, increased with age. In addition, there was a substantial increase in levels of 5’-methyl-thioadenosine (MTA) with age, which can be produced from SAM through the polyamine biosynthesis pathway, and strongly inhibits protein methylation reactions (37). Altered methionine metabolism could impact protein methylation in the aging eye, potentially altering gene expression and signaling pathways.

Next, we asked if some of the changes observed in protein abundance in the aging eye resulted in detectable alterations in the corresponding metabolite pathways. Although many of the cuticle constituents are unlikely to be soluble, we observed significant decrease in one of these metabolites, chitobiose, the major constituent of chitin in the insect cuticle (38). In addition, although glutamine/glutamate metabolism was not significantly enriched in the mummichog analysis, glutamate was identified as a significantly downregulated compound with age (Table S7). In addition, we observed higher levels of the glutamate-related metabolite 2-Pyrrolidone-5-carboxylic acid (pyroglutamic acid) in the aging eye. Together, this suggests, as indicated by the proteomic analysis, the eye experiences alterations in glutamate/glutamine metabolism during aging.

Overall, the metabolomic profiling reveals that there are substantial changes in metabolism in the aging *Drosophila* eye, particularly in pathways associated with folate metabolism. Taken together with the alterations in protein abundance in the aging eye, and integration with transcriptomic analysis in aging eyes and photoreceptors, our data suggest that complex changes in gene expression and metabolism occur in the aging eye.

## Discussion

Several studies have examined proteomic or metabolomic changes in aging *Drosophila* (14, 15, 39–42), but to our knowledge, this is the first study to examine the aging proteome and metabolome specifically in the fly eye. Our data show that the proteomic and metabolomic changes that occur in the aging eye are largely distinct from those that occur in the brain or the body. In the eye, we observed significant age-associated increases in the level of mitochondrial proteins involved in glutamate/glutamine and GABA metabolism, together with decreases in the abundance of proteins that regulate calcium homeostasis, and proteins that are structural constituents of the cuticle and corneal lens. In contrast, proteomic studies in the aging head have reported increased abundance of metabolic proteins involved in oxidative phosphorylation, the TCA cycle, and branched amino acid degradation, together with decreased levels of proteasomal/ubiquitination and ribosomal proteins (14, 15). Moreover, the decline in protein synthesis and increase in proteome variation with age in the head suggest that there is an overall decrease in proteostasis in the aging brain (14, 15), which we did not observe in the eye. In fact, our data show that there is decreased heterogeneity in the aging proteome (Fig. 2B), and that there may be upregulated protein synthesis of at least some mitochondrial proteins in the aging eye because their increase in protein abundance cannot be explained by higher transcript levels. For example, among the glutamate metabolism enzymes that increased with age, only Gs1 was also upregulated at the transcript level in the eye (Fig. 5C). We note that although these proteins are largely localized in mitochondria, they are encoded by the nuclear genome and were all detected in both the eye and photoreceptor nuclear RNA-seq experiments. Overall, the limited similarities between the aging eye and head proteomes were only apparent in comparisons with much older head samples, suggesting that the eye experiences aging more rapidly than the brain.

In contrast to the decrease in variation in the aging proteome in the eye, there was a substantial increase in variation between samples in the aging metabolome (compare Fig. 2B and 6B). This observation suggests that metabolism becomes dysregulated in the aging eye, raising the possibility that altered metabolism could account for the heterogeneity in loss of visual function and retinal degeneration in individual wild-type flies with age. Previous metabolic profiling in whole aging flies identified decreases in pathways associated with glycolysis, fatty acid metabolism, and changes in levels of several amino acids and neurotransmitters including glutamine and GABA (40). In contrast, our metabolite analysis in the eye suggests that the pathways most impacted by aging in this tissue are associated with folate and purine metabolism, with strong downregulation of many pterin-containing metabolites (Fig. 6C, D). Purine and phenylalanine metabolic pathways feed into biosynthesis of pterin derivatives such as biopterin and isoxanthopterin. These pterins are also required for biosynthesis of the red eye pigments in flies, drosopterins (43), and tetrahydropterin, an essential cofactor for several enzymes involved in aromatic amino acid degradation and neurotransmitter biosynthesis. Interestingly, proteins involved in tetrahydropterin biosynthesis are among the few proteins with increased synthesis in the aging head (14). In addition to folate metabolism, we also observed alterations in several metabolites in pathways associated with regeneration of the methyl donor SAM, suggesting that methylation reactions might become dysregulated in the aging eye. In particular, the strong increase in levels of MTA might be expected to have an inhibitory effect on protein methylation reactions (37). Alterations in SAM metabolism and regeneration, particularly in the fat body of *Drosophila*, impact lifespan and are a hallmark of the aging metabolome in the fly body (34, 39, 41, 44). For example, overexpression of the SAM level regulator Gnmt in the fat body of aging flies buffers the age-associated increase in SAM levels and extends lifespan (34). Gnmt is regulated at the transcriptional level by foxo (FBgn0038197), and is upregulated when the Toll pathway is activated (32, 34). The upregulation of several immune response proteins and regulators suggests that there may be a chronic activation of the immune response pathway in the aging eye, which could be responsible for the increase in Gnmt levels and for the alterations in SAM pathways. However, it is unknown whether the increase in Gnmt in aging eyes and photoreceptors impacts SAM levels, function, or protects against retinal degeneration. Moreover, it is unclear whether there are any changes in protein methylation in the aging eye. Notably, Gnmt was one of the few proteins that was highly upregulated in both the eye and brain (Fig. 3C), suggesting that its increased expression is a common feature of aging across multiple tissues in flies.

One of the most intriguing and unexpected observations from the global metabolite profiling was that the levels of riboflavin (Vitamin B2) were significantly decreased in the aging eye (Fig. 6C). Riboflavin cannot be synthesized by animals including *Drosophila*, and knockdown of the sole Riboflavin transporter in flies (Rift; FBgn0039882) decreases mitochondrial activity, alters mitochondrial morphology, and reduces lifespan (45). Moreover, feeding flies with riboflavin extends lifespan, particularly under conditions of constant light exposure (46, 47). Because riboflavin is required for production of FMN and FAD, which are cofactors for many different flavoenzymes involved in metabolism, decreased riboflavin levels could have a wide impact on metabolic pathways in the aging eye. For example, synthesis of pyridoxal phosphate, the biologically active form of vitamin B6 (pyridoxine), also requires FMN (48); thus deficiencies in riboflavin levels could impact many metabolic pathways that require vitamin B derivatives. Notably, while acute short-term depletion of riboflavin in mouse cells results in the degradation of flavoenzymes that lack bound cofactor, longer term riboflavin depletion increased levels of thiamine and pyridoxal bound enzymes, potentially as part of a metabolic stress response (49). We observed pyridoxal phosphate binding as an enriched GO molecular function term in the proteins that increased with age in the eye (Fig. 4A), and pyridoxal phosphate is an essential cofactor for several of the enzymes involved in glutamate metabolism and GABA biosynthesis (Fig. 4C). Thus, our data raise the possibility that decreased levels of riboflavin could be a major contributor to altered metabolism in the aging eye, and might also impact the aging proteome. Riboflavin deficiency in animal models can induce cataracts, and supplementing dietary intake of riboflavin can reduce the risk of some forms of cataract in people (36, 50). Cataracts involve dysfunction of the lens due to damage to its protein components; although the anatomy of the *Drosophila* and vertebrate eye differs substantially, our observation that three of the four major protein components of the corneal lens in the fly eye significantly decrease in abundance with age suggests that there may be some parallels between age-associated dysfunction in the lens of both vertebrates and flies.

The possibility that nutritional deficiencies in aging flies could contribute to dysfunction, paralleling age-associated visual diseases such as cataracts, leads to the question of whether other aspects of the proteomic and metabolomic changes in the aging eye mirror those observed in vertebrate brain and eye. Similar to our observations, global metabolomic profiling in the aging mouse eye revealed substantial changes in metabolite levels particularly in the retina and optic nerve, indicating alteration in pathways affecting mitochondrial metabolism and methylation (51). Based on the observations from both the proteomic and metabolomic profiling of the aging *Drosophila* eye, we propose that the changes in levels of proteins involved in calcium homeostasis and glutamate/glutamine metabolism mimic several aspects of aging in humans, likely more broadly in neurons rather than specifically in the eye, suggesting that the fly eye may provide a good model for these pathways. Calcium homeostasis and signaling are critical for proper neuronal function, and altered calcium levels and signaling occur during aging and can impair cognitive function (52). *Drosophila* photoreceptors, like several other types of neurons, are susceptible to calcium excitotoxicity; prolonged calcium influx resulting from unrestrained phototransduction induces oxidative stress and eventual retinal degeneration (27). We observed calcium binding as one of the enriched GO terms in the proteins that were downregulated with age, and two of the major proteins that buffer calcium levels in photoreceptors are downregulated with age: Calmodulin (Cam) and Cpn. Since maintaining proper calcium homeostasis is critical for photoreceptor survival, and indeed for neuronal health in general, we propose that the decline in levels of these calcium buffering proteins might sensitize the aging eye to retinal degeneration. In addition to dysregulated calcium homeostasis, our data indicate that glutamate/glutamine metabolism and GABA biosynthesis become altered in the aging eye. Alterations in glutamate/glutamine metabolism are implicated in the pathogenesis of human frontotemporal dementia, and involve a complex interplay between these metabolic pathways in neurons and glia (53). While our cell type-specific transcriptome profiling in the eye has previously focused on photoreceptors, only one of the upregulated proteins involved in glutamate/glutamine pathways showed an increase in the transcript level in the eye (Fig. 5C; Gs1). Thus, it will be important to examine other cell types in the aging *Drosophila* eye, particularly those cells such as cone cells that play a glial-like support role for photoreceptors (9). Since *Drosophila* do not show signs of retinal degeneration at day 40 (11), we propose that the alterations in the proteome and metabolome observed in the aging eye in this study precede retinal degeneration, but may provide clues as to the factors that sensitize older flies to photoreceptor loss. These age-regulated pathways may provide targets for neuroprotective approaches to protect aging photoreceptors from damage and retinal degeneration.

## Experimental Procedures

### Fly genetics and aging

*Rh1-Gal4>UAS-GFP^KASH^* flies (*w^1118^;; P{w^+mC^=[UAS-GFP-Msp300KASH}attP2, P{ry^+t7.2^=rh1-GAL4}3, ry^506^*) (11) were raised in 12:12h light:dark cycle at 25°C. Flies were collected on the day of eclosion (day 1), and transferred to fresh vials every two days for aging studies. Eyes (including the lamina) were dissected from adult male flies (2 eyes/fly) using micro-dissection scissors and trimmed to remove excess brain material. Four biological samples were analyzed for each age for both proteomic and metabolomic studies.

### Proteomic sample preparation and proteolytic digestion

Global proteomic mass spectrometry methods were based on previously described approaches (54, 55). Dissected eyes (100/sample) were collected in phosphate-buffered saline (PBS) and flash frozen in liquid N_2_. Samples were homogenized in 25 μL 8 M urea in 100 mM Tris-HCl (pH 8.5) in 1.5 mL tubes using a microtip-based sonication system (BRANSON SONIFER 250, BRANSON SONIC POWER COMPANY, Eagle R. Danbury, CT 06810, USA) at vendor defined “2.5 Output Control” with 20 second bursts for 5 times. The resulting solutions were next subjected to end-over-end rotation at room temperature for 2 hours to enable full dissolution and homogenization of protein, followed by centrifugation at 14,000 rpm for 10 minutes to pellet debris and to retrieve protein extracted solution. Protein concentrations were determined using a Bradford protein assay (BioRad) and colorimeter-EPOCH|2 (BioTek Instruments, Inc., Winooski, VT 05404-0998, U.S.A.) employing vendor provided protocols. Protein samples in equal amounts (20 μg) were next subjected to reduction of Cys-Cys bonds of proteins with 5 mM tris(2-carboxyethyl)phosphine hydrochloride (TCEP) and alkylation with 10 mM chloroacetamide (CAM) to protect the reduced Cys residues from potential recombination reactions. The solutions were then diluted with 100 mM Tris-HCl (pH 8.5) to achieve a final 1.6 M urea concentration. Proteolytic digestion was performed overnight at 37°C using 0.5 μg Trypsin/Lys-C Mix Mass Spectrometry Grade for each sample (Promega Corporation, Madison, WI 53711-5399, U.S.A.). The resulting peptides were de-salted using Sep-Pak^®^ Vac 1cc C18 Cartridges, 50 mg Sorbent per Cartridge, 55-105 μm Particle Size (Waters Corporation Milford, MA 01757, U.S.A.) employing a vacuum manifold (Waters Corporation Milford, MA 01757, U.S.A.). Briefly, columns adapted onto the extraction manifold were first washed two times with 1 mL ACN, and equilibrated three times with 1 mL 0.1% TFA in MS-grade water. Peptides from each digestion solution were then subjected to immobilization on reverse-phase material by a gentle application of vacuum into the extraction manifold vacuum chamber to move each solution three times by collecting the flow-through fractions and transferring them again onto the same column. Next, the peptide-bound reverse-phase columns were washed three times with 1 mL of 0.1% TFA in MS-Grade H_2_O and then eluted by passing 250 μL of ACN/H2O 70/30 (v/v; 0.1% TFA) three times. Elution fractions were combined for subsequent processing, and dried using a speed vacuum system.

### Tandem Mass Tags (TMT) based peptide labelling, reaction quenching, mixing and fractionation

Dried peptide samples were subjected to TMT (Tandem Mass Tags) based labelling using a 10plex kit (TMT10plex™ Isobaric Label Reagent Set, 8 x 0.2 mg). The TMT channels TMT126, TMT127N, TMT127C, and TMT128N, were employed for the labelling of the four “day 10” samples; and the TMT channels TMT128C, TMT129N, TMT129C, and TMT130N were employed for the labelling of the four “day 40” samples. Briefly, each dried sample was reconstituted in 300 μL of 50 mM Triethylammonium bicarbonate (TEAB) and dry labeling reagents were dissolved in 40 μL of acetonitrile (ACN). Reconstituted peptide solutions (100 μL, i.e. 20 μg equivalent protein digest) were then moved to the respective labelling reagent-vials and kept at room temperature for overnight to label the peptides. The labelling reaction was next quenched by adding 8 μL of 5% hydroxylamine at room temperature for > 15 minutes. Labelled peptide solutions were then mixed together and dried using a speed vacuum system. The dried labelled peptide mixture was next fractionated using reversed phase fractionation columns (8 fractions) employing vendor provided protocols (Pierce Biotechnology, Waltham, MA). The resulting 8 fractions were dried using a speed vacuum system and re-constituted in 0.1% formic acid (30 μL) prior to nano-LC-MS/MS analysis, as described below.

### Nano-LC-MS/MS Analysis

Nano-LC-MS/MS analyses were performed on an Orbitrap Fusion™ Lumos™ mass spectrometer (Thermo Fisher Scientific) coupled to an EASY-nLC™ HPLC system (Thermo Scientific). Re-constituted fractionated peptide samples (5 μL eq.) were loaded onto a reversed phase PepMap™ RSLC C18 column (2 μm, 100 Å, 75 μm x 25 cm) with Easy-Spray tip at 750 bar applied maximum pressure. The peptides were eluted using a varying mobile phase (MP) gradient from 94% phase A (FA/H2O 0.1/99.9, v/v) to 28% phase B (FA/ACN 0.4/99.6, v/v) for 160 mins.; from 28% phase B to 35% phase B for 5 mins; from 35% phase B to 50% phase B for 14 mins to ensure elution of all peptides; bringing down the MP-composition to 10% phase B for 1 min at 400 nL/min to bring the MP-composition to higher percentage of phase A. The Nano-LC mobile phase was introduced into the mass spectrometer using an EASY-Spray™ Source (Thermo Scientific™). During peptide elution, the heated capillary temperature was kept at 275°C and ion spray voltage was kept at 2.5 kV. The mass spectrometer method was operated in positive ion mode for 180 minutes having a cycle time of 4 seconds. MS data was acquired using a data-dependent acquisition method that included two “MSn” levels. During the survey MS scan or the MSn level 1, using a wide quadrupole isolation, surveying MS scans were obtained with an Orbitrap resolution of 60 k with vendor defined parameters: m/z scan range, 400-1500; maximum injection time, 50; AGC target, 4E5; micro scans, 1; RF Lens(%), 30; “DataType”, profile; Polarity, Positive with no source fragmentation and to include charge states 2 to 6 for fragmentation. Dynamic exclusion for fragmentation was kept at 60 seconds. During MSn level 2, following vendor defined parameters were assigned to isolate and fragment the selected precursor ions: Isolation mode = Quadrupole; Isolation Offset = Off; Isolation Window = 1; Multi-notch Isolation = False; Scan Range Mode = Auto Normal; First Mass = 100; Activation Type = HCD; Collision Energy Mode = Fixed; Collision Energy (%) = 36 for the 1^st^ replicate and 37 for the second replicate MS analysis; Detector Type = Orbitrap; Orbitrap Resolution = 50k; Data type = Centroid; Polarity = Positive; Source Fragmentation = False. The data were recorded using Thermo Scientific Xcalibur (4.1.31.9) software (Thermo Fisher Scientific Inc.).

### Multiplexed proteomic data analysis

The resulting RAW files were analyzed using Proteome Discover 2.2.0.388 (ThermoScientific). A specific TMT 8plex quantification method was formulated using the default TMT 10plex method using the tools available in Proteome Discover 2.2. The MS/MS spectra were searched against *in silico* tryptic digest of a Drosophila protein database (FASTA format) downloaded from the UniProt sequence database (v. Mar 2018) using the SEQUEST HT search engine that also accounted for “common contaminants”. In order to carry out the search, following specific search parameters were applied to vender provided “processing” and “consensus” workflow templates that correspond to Thermo “Fusion” instruments: Trypsin as the proteolytic enzyme; searched for peptides with a maximum number of 2 missed cleavages; precursor mass tolerance of 10 ppm; and a fragment mass tolerance of 0.6 Da. Static modifications used for the search were: 1) carbamidomethylation on cysteine(C) residues; 2) TMT 6plex label on lysine (K) residues and the N-termini of peptides. Dynamic modifications used for the search were oxidation of methionine, and acetylation of N-termini. Percolator False Discovery Rate was set to a strict setting of 0.01 and a relaxed setting of 0.05. Values from both unique and razor peptides were used for quantification. In order to account for procedural errors, reporter ion-based responses for proteins were normalized in Proteome Discover 2.2 (Thermo Scientific) using the “total peptide amount” option. Resulting “grouped” abundance values for each sample type; “abundance ratio” values; and respective “p-values (ANOVA)” from Proteome Discover were used to identify statistically increased or decreased proteins for global proteomic comparison between day 10 and day 40 samples.

### Metabolomic sample preparation and HPLC-MS analysis

Dissected eyes (100/sample) were collected in 200 μL methanol in Precellys tubes (Cayman Chemicals #10011152) and flash frozen in liquid N_2_. Protein removal and sample extraction were performed using the Bligh-Dyer extraction protocol, as detailed by Setyabrata et al. (56). Non-targeted metabolite profiling was performed utilizing HPLC-MS, with a reversed-phase C18 chromatographic separation followed by high mass accuracy time-of-flight mass spectrometry, scanning between 100 – 1100 Daltons, as detailed in (56). Compound identification was aided by performing data dependent MS/MS collection on composite samples. Peak deconvolution and integration were performed using Agilent ProFinder (v. B.10) and manually edited for accuracy. Bioinformatics were performed using Agilent’s Mass Profile Professional (v. 13.1). Chromatographic peaks were aligned across all samples. Peak areas were normalized by converting to log_2_ values and applying a 75% percentile shift. Significance analysis was performed using an unpaired t-test with Benjamini-Hochberg FDR correction. Metabolites with p < 0.01 and fold change > 2 were considered significant. Peak annotations were performed using the METLIN (metlin.scripps.edu) metabolite databases, with a mass error of less than 10 ppm. MetaboAnalyst 4.0 (https://www.metaboanalyst.ca/) was also utilized for comparative statistical and pathway analysis.

## Conflict of Interest

The authors declare that they have no competing interests.

## Data Availability

The mass spectrometry proteomics data have been deposited to the ProteomeXchange Consortium via the PRIDE (57) partner repository with the dataset identifier PXD023275 and 10.6019/PXD023275. Reviewers can access these data using: Username: Password: ftalF5Kq. Previously published RNA-seq expression data are accessible through Gene Expression Omnibus (GEO) repository under series accession numbers GSE83431 and GSE106652.

## Acknowledgements

Information from FlyBase was used in this study. The authors thank Purdue University’s Metabolite Profiling Facility for assistance with mass spectrometric analysis for the metabolomic studies. The mass spectrometry proteomic studies performed in this work was performed by the Indiana University Proteomics Core. Acquisition of the IUSM Proteomics core instrumentation used for this project was provided by the Indiana University Precision Health Initiative.

## Funding

Support from the American Cancer Society Institutional Research Grant (IRG #58-006-53) to the Purdue University Center for Cancer Research is gratefully acknowledged. The proteomics work was supported, in part, with support from the Indiana Clinical and Translational Sciences Institute funded, in part by Award Number UL1TR002529 from the National Institutes of Health, National Center for Advancing Translational Sciences, Clinical and Translational Sciences Award. Research reported in this publication was supported by the National Eye Institute of the NIH under Award Number R01EY024905 to VW.

## Notes

### Competing Interest Statement

The authors have declared no competing interest.

